# NetMHCpan 4.0: Improved peptide-MHC class I interaction predictions integrating eluted ligand and peptide binding affinity data

**DOI:** 10.1101/149518

**Authors:** Vanessa Jurtz, Sinu Paul, Massimo Andreatta, Paolo Marcatili, Bjoern Peters, Morten Nielsen

## Abstract

Cytotoxic T cells are of central importance in the immune system’s response to disease. They recognize defective cells by binding to peptides presented on the cell surface by MHC (major histocompatibility complex) class I molecules. Peptide binding to MHC molecules is the single most selective step in the antigen presentation pathway. On the quest for T cell epitopes, the prediction of peptide binding to MHC molecules has therefore attracted large attention.

In the past, predictors of peptide-MHC interaction have in most cases been trained on binding affinity data. Recently an increasing amount of MHC presented peptides identified by mass spectrometry has been published containing information about peptide processing steps in the presentation pathway and the length distribution of naturally presented peptides. Here, we present NetMHCpan-4.0, a method trained on both binding affinity and eluted ligand data leveraging the information from both data types. Large-scale benchmarking of the method demonstrates an increased predictive performance compared to state-of-the-art when it comes to identification of naturally processed ligands, cancer neoantigens, and T cell epitopes.

## Introduction

Cytotoxic T cells play a central role in the immune regulation of pathogenesis and malignancy. They perform the task of scrutinizing the surface of cells for the non-self peptides presented in complex with MHC (major histocompatibility complex) molecules. In cases such peptides are recognized, an immune response can be initiated potentially leading to killing of the infected (mal-functioning) cell. The most selective step in the pathway leading to this peptide presentation is the binding to MHC.

Over the last decades, large efforts have been dedicated to the development of computational methods capable of accurately predicting this event. The accuracy of these methods has improved substantially over the last years, and most recent benchmark results demonstrate that more than 90% of naturally presented MHC ligands are identified at an impressive specificity of 98% (1). This gain in performance is achieved partly by the extended experimental binding data sets made available in the IEDB (2), and partly by the development of novel machine-learning algorithms capable of capturing the information in the experimental binding data in a more effective manner. One such novel method is NNAlign-2.0, allowing the integration of peptides of variable length into the machine-learning framework (3). This novel training approach allows both the incorporation of a larger set of training data, but also and maybe more importantly enables the method to directly learn the length preference presented peptides for each MHC molecule from the experimental binding data (4). Even though most presented MHC class I ligands are of length 9 amino acids, the ability to incorporate length preferences directly into the model is critical as experimental data demonstrate that the length profiles of presented ligands can vary substantially between MHC molecules; prominent examples are the mouse H-2-Kb, with a preference for eight amino acids-long peptides (5) and HLA-A*01:01, where close to one third of MHC presented peptides have a length longer than nine amino acids (6).

Some of the most well documented and applied of methods for predicting peptide binding to MHC class I include NetMHC (4,7), and NetMHCpan (1,8). These tools have over the last years gained increasing interest due to the recent focus on neoantigen identification within the field of personalized immunotherapy (9,10). However, as underlined in several studies including the recent Nature Biotechnology Editorial (11), “neoantigen discovery and validation remains a daunting problem”, mostly due to the relative high false positive rate of predicted epitopes.

One potential cause for this relatively high rate of false positive epitope predictions is the fact that most methods are trained on binding affinity data, and as a consequence only model the single event of peptide-MHC binding. As stated above this binding to MHC is the most selective step in peptide antigen presentation. However, other factors including antigen processing (12) and the stability of the peptide:MHC complex (13) could influence the likelihood of a given peptide to be presented as an MHC ligand. Similarly, the length distribution of peptides available for binding to MHC molecules is impacted by other steps in the processing and presentation pathway, such as TAP transport and ERAP trimming, which are not reflected in binding data in itself (6). Advances in mass spectrometry (MS) have allowed the field of MS peptidomics to move forward. In this context, recent studies (14,15,16) have suggested that training prediction methods on such data rather than binding affinity data could improve the ability to accurately identify MHC ligands. As such, MS peptidome data would contain the comprehensive signal of antigen processing and presentation rather than just MHC binding affinity. Moreover, MS peptidome data generated by immunopeptidomic studies would contain precise information about the allele-specific peptide length profile preferences not available in the MHC binding affinity data sets.

Identification of MHC bound peptides by mass spectrometry thus holds great promise for the generation of large scale data sets characterizing the peptidome specific for individual MHC molecules (15,17), and potentially also for the identification of T cell epitopes (18). It is however clear that, within the foreseeable future, the number of MHC molecules characterized by such MS studies will remain limited. In this context, large efforts have over the last decades been dedicated to experimentally characterize the peptide binding space of MHC molecules using semi high-throughput MHC-peptide binding affinity assays (19,20), enabling binding specificity characterization of a large set of MHC molecules from different species.

The IEDB contains a comprehensive set of MHC binding and ligand data available in the public domain. While this data set contains binding affinity data characterizing more than 150 different MHC class I molecules (from human, non-human primates, mouse, and life-stock), at the onset of this study only 55 MHC class I molecules were characterized by MS peptidome data. This imbalance made us suggest a novel machine learning approach integrating information from both types of data (binding affinity and MS ligands) into a combined framework benefitting from information from the two worlds. The proposed framework is “pan-specific” as it can leverage information across MHC molecules, data types, and peptide lengths into one single model. We hence expect this approach to achieve superior predictive performance compared to models trained on the two data types individually, and also achieve an improved performance when it comes to predicting length profile preferences of different MHC molecules.

While recent works have demonstrated the improved ability to identify MHC ligands using methods trained on MS peptidome data (14,15), limited data is available on their impact for the identification of T cell epitopes. In this work, we focus on demonstrating the improved prediction performance not only on large sets of MS peptidome data but also on T cell epitope data independent from the data used to train the new predictor.

## Materials and Methods

### Data sets

Data on all class I MHC ligand elution assays available in IEDB database (www.iedb.org) were collected including the ligand sequence, details of the source protein, position of the ligand in the source protein and the restricting allele of the ligand. There were 160,527 distinct assays in total and the length of the ligands ranged from 4-37. All lengths with a count of ligands at least 0.5% of total ligands were selected for further analysis which included lengths 8-15 and comprised of 99% of the assay entries.

The restricting MHC molecule of the ligands were analyzed and entries with alleles listed unambiguously were selected. For example, some entries where the HLA alleles are listed as just the gene name and alleles from chicken, horse, cow and mouse for which we did not have binding prediction algorithms were excluded. Representative alleles were assigned for entries where only supertypes were listed (e.g. HLA-A*26:01 for HLA-A26). Thus there were 127 class I molecules from human and mouse in the selected data set. Redundant entries with same ligand sequence and MHC molecule were removed and MHC molecules with at least 50 ligand entries were selected. This included 55 class I molecules and the number of available ligands per molecule varied widely from 50 to 9500.

We hypothesized that some of the ligands could be artefacts of the elution assays and therefore their source proteins could be false positive as antigens. A protocol was designed to identify such false positive antigens and exclude them from the final data selected. The protocol identified proteins that had significantly lesser number of predicted binders among ligands than expected of random peptides using binomial probability distribution. Five sets of random peptides were generated from the ligand sequences by shuffling the amino acid residues within the ligands. Binding affinity was then predicted for the original ligands and random peptide sets for their corresponding alleles. The median of the predicted percentile ranks of the five random sets was estimated and assigned as the binding affinity of the random peptides. Based on a predicted binding affinity cut-off of percentile rank 1.0, the number of predicted binders among the original ligands and the random peptide sets were estimated. Five proteins were thus identified as false positives and ligand entries from these proteins were excluded from the data set.

The final data set had 85,217 entries in total with ligand length ranging from 8 to 15. The ligands originated from 14,797 source antigens and were restricted by 55 unique HLA molecules.

Random artificial negatives were generated for each MHC molecule covered by eluted ligand data by sampling randomly 10*N peptides of each length 8-15 amino acids from the antigen source protein sequences, where N is the number of 9mer ligands for the given MHC molecule.

### Neural network training

The NNAlign training approach with insertions and deletions (3) was extended by adding a second output neuron as shown in figure 1. This was done to allow combined training on binding affinity and MS eluted ligand data. Binding affinity values are measured as IC50 values in nM (aff) and can be rescaled to the interval [0,1] by applying 1-log(aff)/log(50,000), representing continuous target values (21). For eluted ligands the strength of the interaction between peptide and MHC molecules is unknown, therefore a target value of 1 is assigned to binders and 0 to artificial negative peptides (see above).

**Figure 1:**
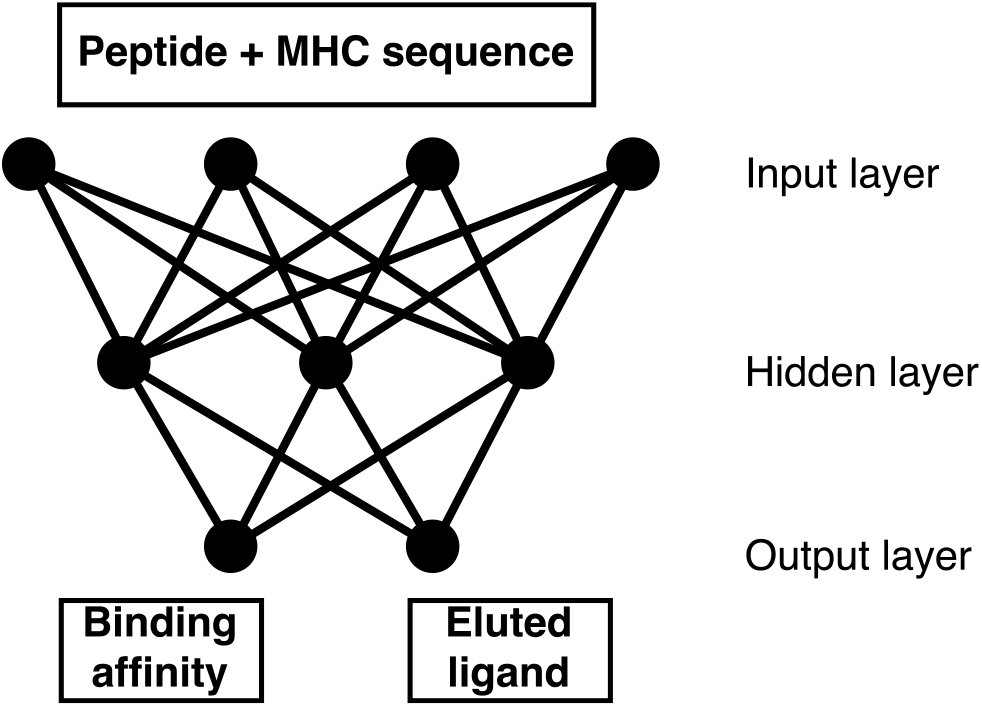
Visualization of the neural networks with two output neurons used for combined training on binding affinity and eluted ligand data.

In this network architecture weights between the input and hidden layer are shared between the two input types (binding affinity/ligand), and weights connecting the hidden and output layer are specific for each input type. During neural network an example is randomly selected from either data set and submitted to forward-and backpropagation according to the NNAlign algorithm (3). In this setting, we define one training epoch as the average number of iterations needed to process each data point in the smaller data once.

A neural network ensemble was trained as described by Andreatta et al. (1) using 5-fold nested cross-validation. Networks with 60 and 70 hidden neurons were trained leading to an ensemble of 40 networks in total.

The inputs to the neural networks consisted of the peptide and the MHC molecule in terms of a pseudo sequence (8). All peptides were represented as 9-mer binding cores by the use of insertions and deletions as described by Andreatta et al. (4) and encoded using BLOSUM encoding (21). As in the earlier work by Andreatta et al. (4), additional features for the encoding of peptides included: the length of the deletion/insertion; the length of peptide flanking regions, which are larger than zero in the case of a predicted extension of the peptide outside either terminus of the binding groove; and the length L of the peptide, encoded with four input neurons corresponding to the four cases L<=8, L=9, L=10, L>=11.

### Performance

In order to benchmark the combined training method described above (referred to as BA+EL), additional methods with only one output but otherwise identical setup were trained on binding affinity data only (BA data) and eluted ligand data only (EL method). Performance was measured as area under the receiver operating curve (AUC), a value of AUC=0.5 indicates random model performance while an AUC=1 represents a perfect model. AUC values were calculated for each MHC allele separately and subsequently binomial tests were performed to compare the different models.

### Length preference of MHC molecules

For all MHC molecules shared between the binding affinity and eluted ligand data sets, we generated predictions for 80,000 random natural peptides of lengths 8-15 amino acids (10,000 of each length). From the top 2% predictions, the frequency of each peptide length was estimated. Subsequently Pearson's correlation coefficient was calculated between the frequencies observed in the eluted ligand data set and the frequencies predicted by 4 models (BA only, EL only, binding affinity of BA+EL, and eluted ligand predictions of BA+EL)

### Leave-one-out validation

Leave-one-out experiments were performed for all MHC molecules present in the eluted ligand data set. For this, a given MHC molecule was removed from the eluted ligand data set, then the BA+EL method was trained in five-fold cross-validation as described above, omitting multiple random initializations, resulting in an ensemble of 10 networks. Performance of the leave-one-out models is compared to an ensemble of neural networks of the same size trained on the complete data set. Further predictions are made for 80,000 peptides of lengths 8-15 amino acids derived from natural proteins to evaluate a model’s ability to predict the length preference of an MHC allele that was not part of the eluted ligand training data.

### The final NetMHCpan-4.0 method implementation

The final neural network ensemble of the NetMHCpan-4.0 method is trained on binding affinity and eluted ligand data as described above using 5-fold cross-validation. Networks with 56 and 66 hidden neurons (in accordance with earlier NetMHCpan implementations) were trained using 10 distinct random initial configurations, leading to an ensemble of 100 networks in total.

Percentile rank scores was estimated from predicted EL and BA binding values from a set of 125,000 8-12mer random natural peptides (25,000 of each length)

### Validation on external data sets

A dataset of eluted ligands was obtained from Pearson et al. (17). Also, a set of positive CD8 epitopes was downloaded from the IEDB. The epitope set was identified using the following search criteria “T cell assays: IFNg”, #x201C;positive assays only”, “MHC restriction Type: Class I”. Only entries with fully typed HLA restriction, peptides length in the range 8-14 amino acids, and with annotated source protein sequence were included. Positive entries with a predicted rank score greater than 10% using NetMHCpan-3.0 were excluded to filter out likely noise (6). For both the T-cell epitope and eluted ligand data sets, negative peptides were obtained by extracting all 8-14mer peptides from the source proteins of the eluted ligands and subsequently excluding peptides-MHC combination found with an exact match in the training data (both binding affinity and eluted ligand data sets). The final eluted data set contained 15,965 positive ligands restricted to 27 different HLA molecules, and the IEDB T cell epitope data set 1,251 positive T cell epitopes restricted to 80 HLA molecules.

A Frank value was calculated for each positive-HLA pair as the ratio between the number of peptides with a prediction score higher than the positive peptide and the number of peptides contained within the source protein. The Frank value is hence 0 if the positive peptide has the highest prediction value of all peptides within the source protein, and a value of 0.5 in cases where an equal amount of peptides has a higher and lower prediction value compared to the positive peptide.

An unfiltered eluted ligand data set was obtained from Bassani-Sternberg et al. (22). This data sets consisted of eluted ligand data from 6 cell lines each with fully typed HLA-A, B and C alleles. A data set was constructed for each cell line, including all 8-13mer ligand as positives, and 5 times the total number of ligands random natural negatives for each length 8-13. That is if a data set contained 5,000 ligands, 5*5000 = 25,000 random natural peptides of length 8, 9, 10, 11, 12, and 13 were added as negatives arriving at a final data set with 155,000 (5000 + 6*25000) peptides.

## Results

We trained the NetMHCpan method version 4.0 for the prediction of the interaction of peptides with MHC class I molecules integrating binding affinity and MS eluted ligand data. Combined training was achieved by adding a second output neuron to the NNAlign approach described previously (3). In this setup, the first output neuron returns a score of binding affinity, and the second output neuron a score of ligand elution. As described in materials and methods, the model parameters between the input and hidden layer of the artificial neural network are shared between the two input types. Thanks to this network architecture, we expect the model to be able to combine informative patterns found in the two data types, boosting performance for both output neurons. To demonstrate this, we compared the performance of the BA+EL method to the BA method, trained only on binding affinity data and the EL method trained only on eluted ligand data. Figure 2 shows the mean performance per MHC allele of the four methods on four different data sets given in in terms of AUC (for details see Supplementary Table 1). From this analysis, it is clear that especially the BA+EL method with EL predictions performs much better on binding affinity data than the EL only method. This observation strongly suggests that the EL only method, as a results of the small number of only 55 different MHC molecules included in the eluted ligand data set, has limited pan-specific potential compared to the BA+EL EL method trained on data from 169 MHC molecules included in the combined binding and MS eluted ligand data set.

**Figure 2:**
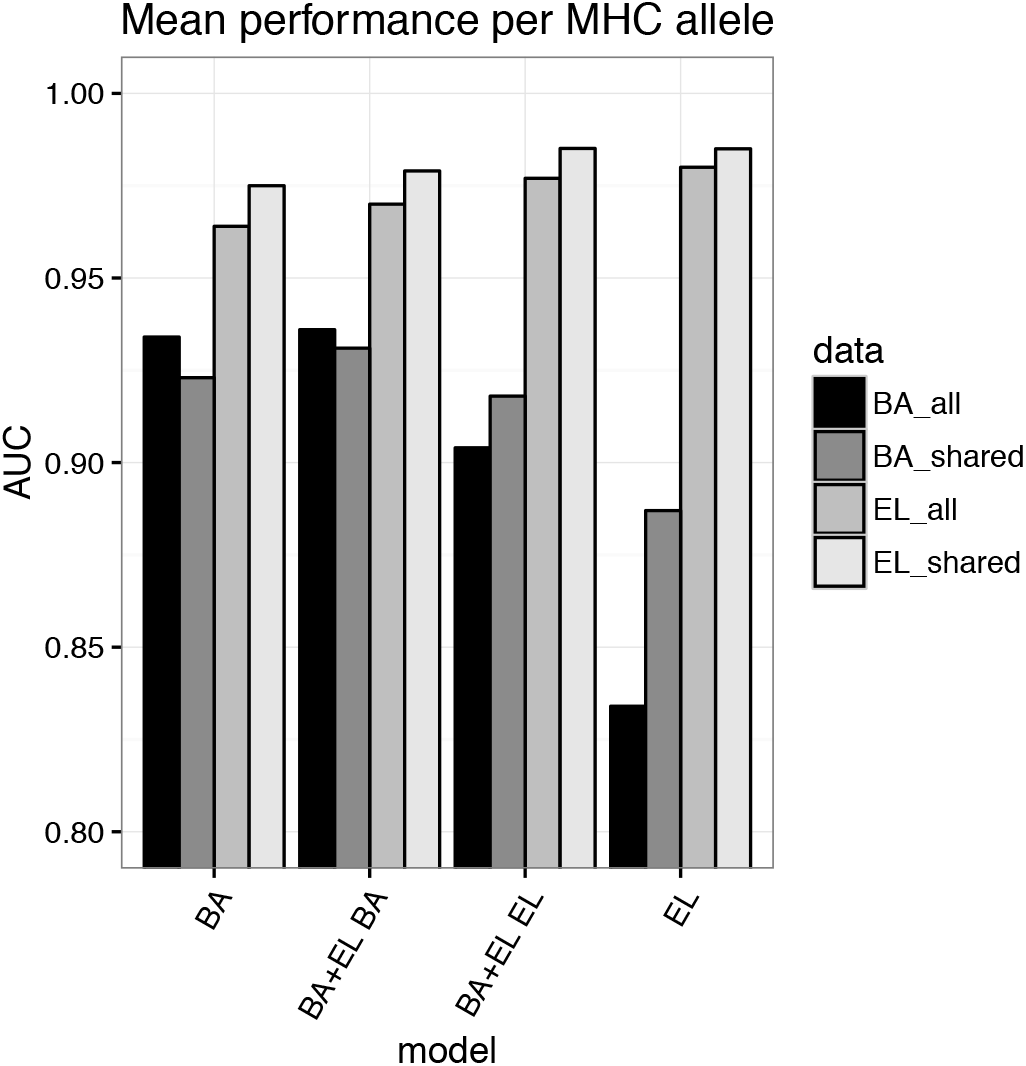
Mean performance per MHC molecule measured in terms of AUC for the four methods; BA (trained on binding affinity data only), EL (trained on eluted ligand data only), BA+EL BA (the binding affinity prediction value of the model trained on the combined binding affinity and eluted ligand data), and BA+EL EL (the eluted ligand likelihood prediction value of the model trained on the combined binding affinity and eluted ligand data) The methods were evaluated on all binding affinity (all_BA) data and all eluted ligand (all_EL) data including negative peptides derived from source proteins, and on data sets restricted to alleles occurring in both binding affinity and eluted ligand data sets (shared_BA, and shared_EL).

### Peptide length preference of MHC molecules

We next set out to investigate how well the different methods could capture the peptide length preferences of individual MHC molecules. For this, we predicted binding scores for a set of random natural peptides of lengths 8-15 amino acids and calculated the frequencies of peptides of different lengths in the top 2% of predictions. In figure 3a-c, we visualize examples of such peptide length preference profiles predicted by the BA, BA+EL BA, BA+EL EL, and EL only methods. The depicted MHC molecules are known to have preferences for different peptide lengths. All HLAs have a preference for 9mer peptides. However, HLA-A*01:01 has an increased preference for 10-mers compared to average, HLA-A*02:01 has a strong preference for 9-mers only, and HLA-B*51:01 has an increased preference for 8-mers compared to average (6). Binding affinity predictors often overestimate the amount of binding 10-mer peptides due to their over-representation in the binding affinity data set (4), which is also visualized in figure 3.

**Figure 3:**
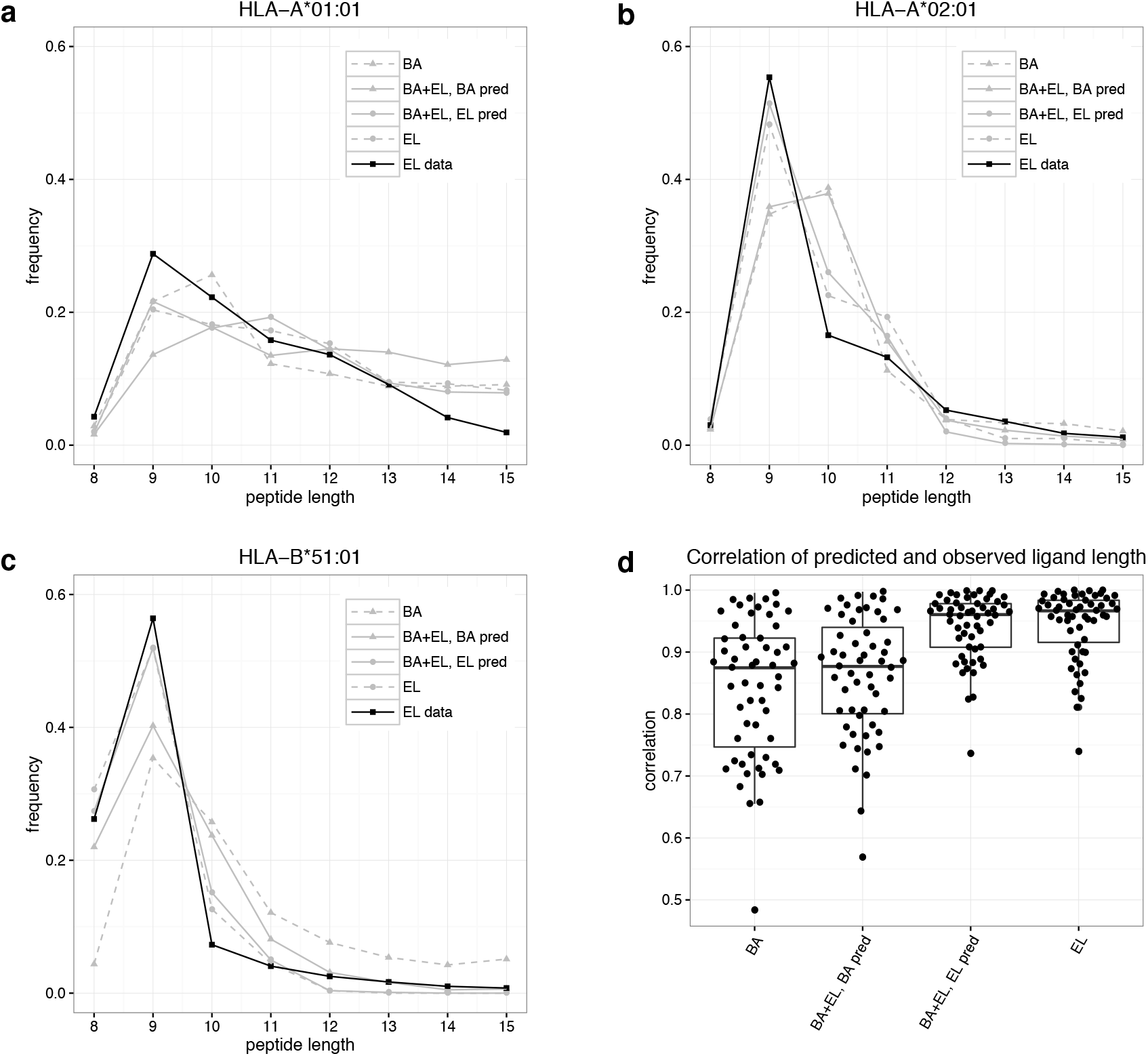
**a-c)** Predicted length preference of selected MHC molecules according to different models. Binding to selected HLA molecules was predicted for 80,000 8-15-mer peptides and the frequency of peptide lengths in the top 2% predicted peptides calculated. **d)** Correlation of predicted and observed ligand length for different models. Binding to all HLA alleles present in both binding affinity and eluted ligand data sets was predicted using the four different prediction methods for 80,000 8-15-mer peptides. Subsequently, the occurrence of different peptide lengths in the top 2% predicted peptides for each molecule was calculated, and the correlation coefficient between these frequencies and the length frequencies in the eluted ligand data set calculated.

Next, we extended the analysis to all MHC molecules included in the eluted ligand data set, calculating the correlation between observed and predicted length frequencies for each prediction method. This analysis (figure 3d) clearly confirms the results obtained from the 3 case examples, namely that the two methods BA+EL EL and EL only show significantly higher power for predicting the peptide length preference of individual MHC molecules compared to the two methods trained to predict binding affinity (BA, and BA+EL BA).

The predictions for the two eluted ligand likelihood models only show low performance for one molecule; HLA-B41:04. This molecule is however only characterized by 52 eluted ligands, whose length profile forms an unusual bimodal distribution with peaks at length 9 and 11 (data not shown).

### Leave-one-out experiments on eluted ligand data

In the above experiment, the MHC molecules used for the peptide length preference evaluation were also included as training data of the EL prediction methods. This naturally leads to a bias in the performance evaluation. To address this, and to access the pan-specific potential of the BA+EL EL prediction method, we conducted a leave-one-out experiment. Here, a given MHC molecule was removed from the eluted ligand data set, and the BA+EL method retrained as described in material and methods. Next, both the predictive performance (estimated in terms of AUC for separating the known ligands from the artificial negatives) and the ability to predict the peptide length preference were evaluated. The result of the benchmark is shown in figure 4.

**Figure 4:**
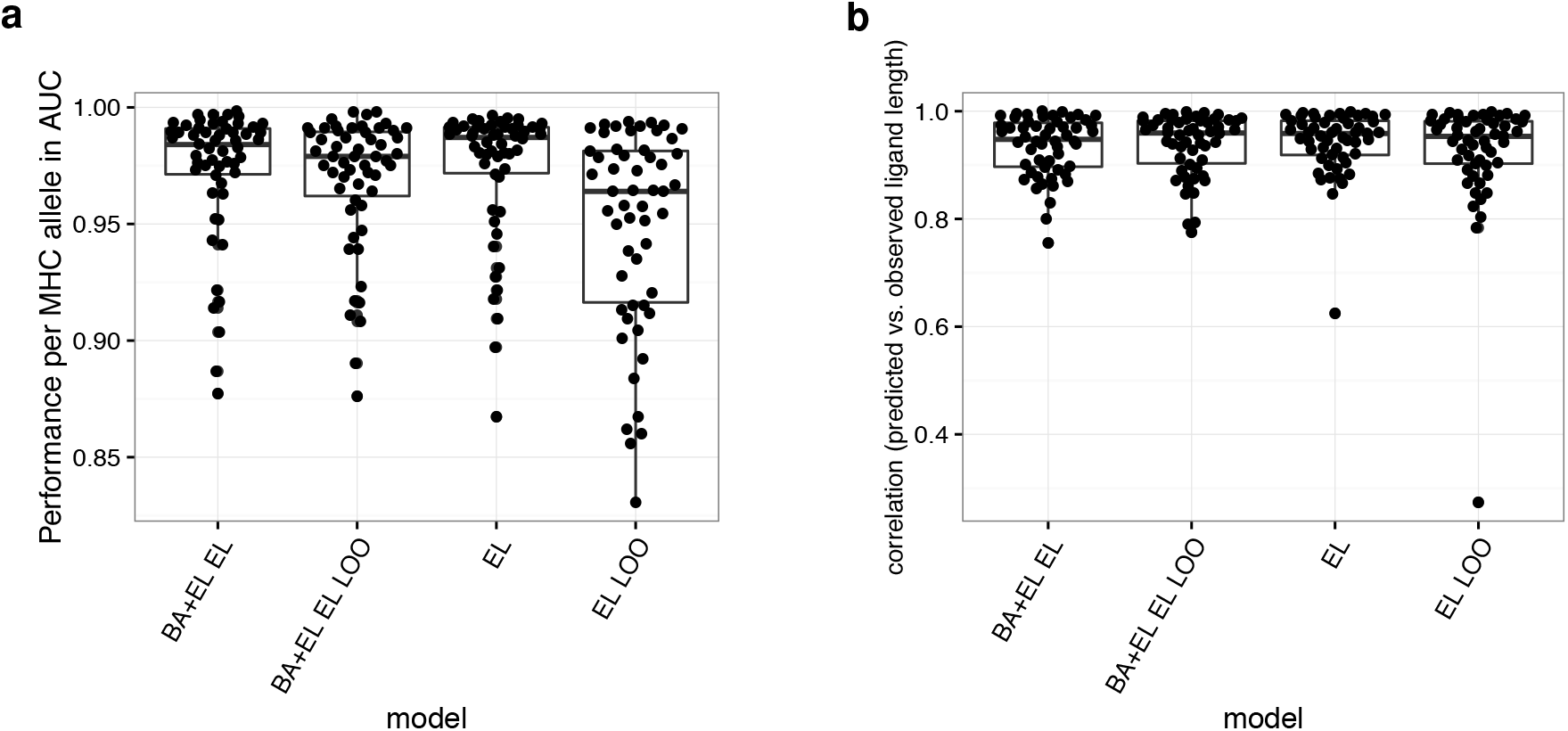
Eluted ligand leave-one-out experiments. **a)** Performance per MHC allele of a model trained on all data and a model where the eluted ligand data of a given allele was left out of the training process. **b)** Correlation of predicted and observed ligand length for a model trained on all data and the leave-one-out models.

This figure clearly confirms the pan-specific power of the BA+EL method. In terms of the predictive performance (figure 4a), the LOO methods display, as expected, a slight decrease in performance compared to a method trained and evaluated on all data (the all data method). When looking at the performance for predicting the peptide length profile (figure 4b), the LOO methods display a very high performance. Only in one case, the EL LOO method shows a substantial drop in performance for the left out MHC molecule. This case is H2-Kb, the only mouse molecule in the MS ligand data set with a strong preference for 8mer ligands. The BA+EL EL LOO method is able to predict the length profile of H2-Kb due to the H2-Kb affinity data present in the BA training data set.

### The NetMHCpan-4.0 method

Having demonstrated the increased predictive power of the BA+EL method when it comes to prediction of peptide binding affinity (the BA+EL BA model), likelihood of being an eluted ligand (BA+EL EL model), and the ability of capturing the MHC specific peptide length binding preferences (also the BA+EL EL model), we set out to construct the final NetMHCpan-4.0 method. This method was trained as the BA+EL method, using 5 fold cross-validation as described in materials and methods. The method is accessible at www.cbs.dtu.dk/services/NetMHCpan-4.0. The functionality is identical to the earlier NetMHCpan implementations with the important additional functionality that two different output options (binding affinity and eluted ligand likelihood) are available. By default, the program returns eluted ligand likelihood scores. An example of the output of the method is shown in Supplementary figure 1.

### Validation on external data sets

The performance of the updated NetMHCpan method was assessed on two independent external data sets; one consisting of 15,965 eluted ligands covering 27 HLA molecules, and another consisting of 1,251 validated CTL epitopes covering 80 HLA molecules reported in the IEDB. The validation data sets were constructed as described in materials and methods. The source protein sequence was identified for each ligand/epitope, and all overlapping 8-14 mer peptides except the ligand/epitope were annotated as negatives. All data points included in the binding affinity and eluted ligand training data sets were excluded from the validation data set. A Frank value was calculated for each positive-HLA pair as described in materials and methods as the ratio of the number of peptides with a prediction score higher than the positive peptide to the number of peptides contained within the source protein. In this manner, we can construct the sensitivity curves presented in figure 5. Two observations are striking from these results. First and foremost, the results clearly demonstrated the increased predictive power of integrating eluted ligand data into the training data of NetMHCpan. In the left panel (the analysis of the eluted ligand data), we can observe that the gain in sensitivity at a Frank threshold of 1% for the EL models (NetMHCpan-4.0 EL or EL only) compared to NetMHCpan-3.0 is 10% (95% versus 85%), and 15% at a Frank threshold of 0.5% (90% versus 75%). These numbers mean that a ligand will have a prediction score within the top 0.5% of its source protein peptides in 90% of the cases using the EL models, compared to only 75% using NetMHCpan-3.0. The results shown in the left panel of figure 5 however also suggest that the two EL models achieve very similar predictive performance when it come to identification of eluted ligands. This is in strong contrast to the results obtained from the IEDB epitope data set (figure 5, right panel). Here, only the NetMHCpan-4.0 EL model demonstrates an improved predictive performance compared to NetMHCpan-3.0.

**Figure 5:**
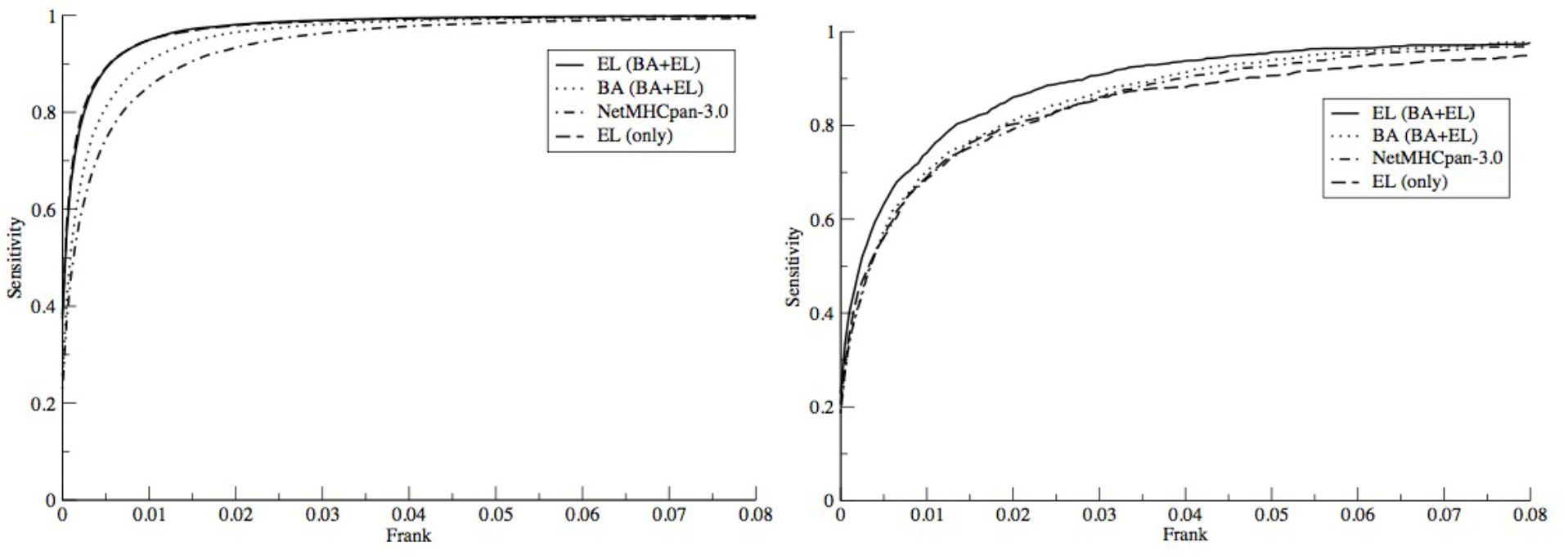
Sensitivity of different models as a function of the Frank threshold on **a)** eluted ligands published by Pearson et al. (17) and **b)** T-cell epitope data downloaded from IEDB.

There are several potential explanations for the improved performance of the EL models on the eluted ligand evaluation data including i) a bias against cysteins specific for the eluted ligand training and evaluation data, ii) as suggested earlier (15) differences in the MHC binding motifs contained within the eluted ligand and *in-vitro* binding data, and iii) the improved prediction accuracy of the ligand length preference (see figure 3d). To investigate i) we repeated the experiment displayed in figure 5, removing all peptides containing one of more cysteins. If the bias against cysteins in the eluted ligand data had any impact on the predictive performance of the proposed method, the bias would be reflected in an altered predictive performance on the reduced data sets. This turned out not to be the case (data not shown) hence suggesting that cysteine bias is not the influencing the relative predictive performance of the different methods. Looking into the differences in the binding motif derived from binding affinity and eluted ligand data respectively for specific HLA molecules, we find differences for most MHC molecules. A few examples are shown in figure 6.

**Figure 6:**
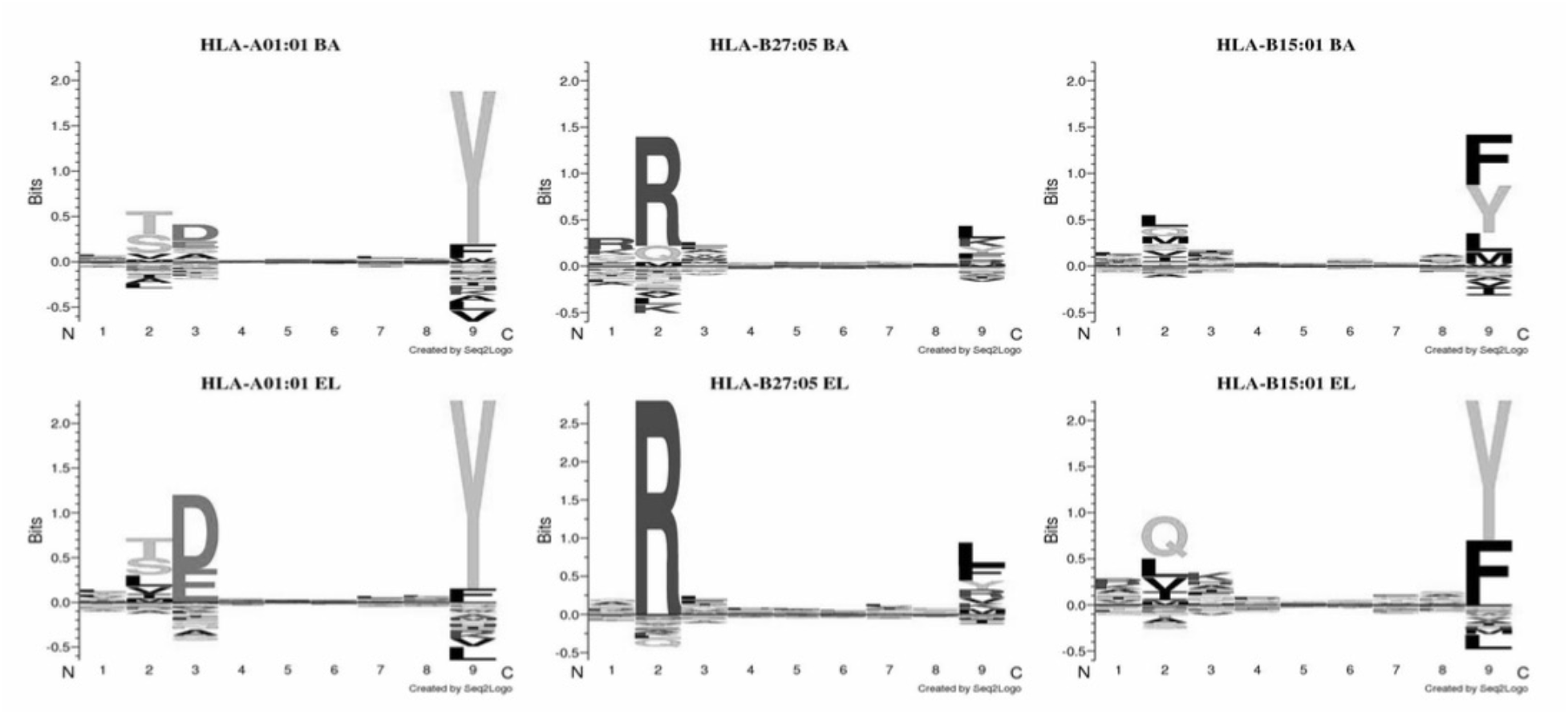
Binding motifs for HLA molecules derived from (upper panel) in-vitro binding affinity data using a binding threshold of 500 nM, (lower panel) eluted ligand data. Logos were made using Seq2Logo with default parameters (30).

These results demonstrated that eluted ligands tend to share more conserved anchor motifs compared to affinity-defined binders. This observation is in agreement with earlier findings suggesting eluted ligands to be more stably bound to MHC-I molecules compared with other affinity matched peptides (13,23). In summary, these analyses suggest that the gained predictive performance of the EL method on the eluted ligand evaluation data is driven by at least two factors; differences in binding preferences between eluted ligand and affinity-defined peptide binders, and the improved prediction accuracy of ligand length preference of the EL methods.

### To be or not to be a ligand

We investigated what prediction threshold to use to best separate ligand from non-ligand peptides. Earlier work by others and us suggests that different MHC molecules present peptides at different predicted binding affinity thresholds (1,24). Given this, it was interesting to investigate to what degree a similar observation could be made for the eluted ligand likelihood predictions produced by the NetMHCpan-4.0 method. To address the question, we compared the predicted ligand likelihood scores of all 15,965 ligands in the Pearson data set. The result of this analysis is displayed as box-plots in the left panel of figure 7.

**Figure 7:**
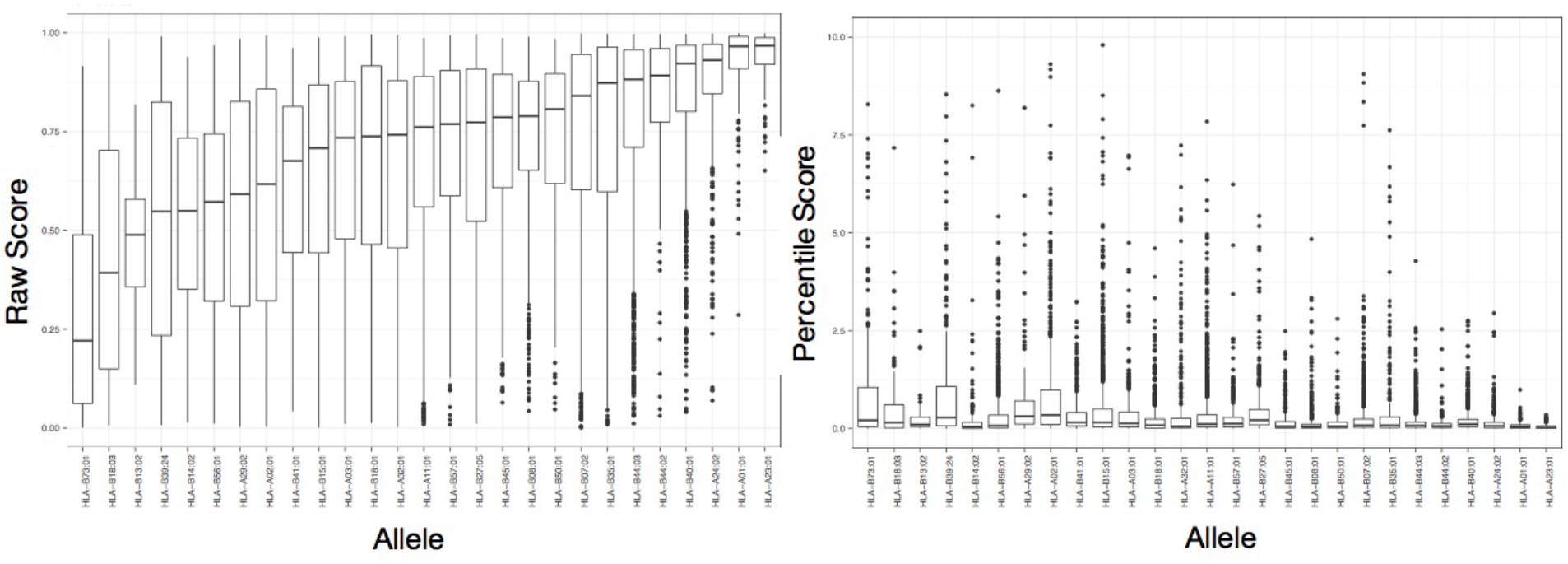
Motivation for using percentile rank score predictions. Box-plot representation of prediction values for the ligands in the Pearson data set. Left panel: Eluted ligand likelihood prediction scores. Right panel: Percentile rank values.

This figure reveals that the likelihood prediction scores for known ligands come out very different for different HLA molecules. The large difference in prediction values between HLA molecules can to a high degree be linked to their absence from the eluted ligand training data. The molecules with lowest median eluted ligand likelihood scores in this figure are molecules absent from the eluted ligand training data set. However, as demonstrated in figure 4 and 5, the fact that an HLA molecule has not been characterized with eluted ligand training data does not impair its predictability. Given this, a natural measure to correct for this great imbalance in prediction score is use percentile rank scores to reconcile and make prediction score comparable between different MHC molecules. The right panel of figure 7 shows the results of such a transformation. Here, eluted ligand likelihood prediction values for each ligand in the Pearson data are transformed to percentile rank scores, and the score distribution is visualized as box plots for each HLA molecule. Given that percentile rank values fall in the range 0-100%, it is apparent that transforming the prediction values into such rank scores, allows for a direct score comparison between HLA molecules.

In light of these results, we next investigated what percentile rank threshold to apply to optimally identify MHC ligands. We assess this by calculating sensitivity/specificity curves as a function of the percentile rank score threshold for a balanced set (max 100 ligands per HLA) of eluted ligands and source protein negatives from the Pearson evaluation data set. The results are shown in figure 8 and confirm earlier findings that the vast majority (96.5%) of natural ligands are identified at a very high specificity (98.5) using a percentile rank threshold of 2%.

**Figure 8:**
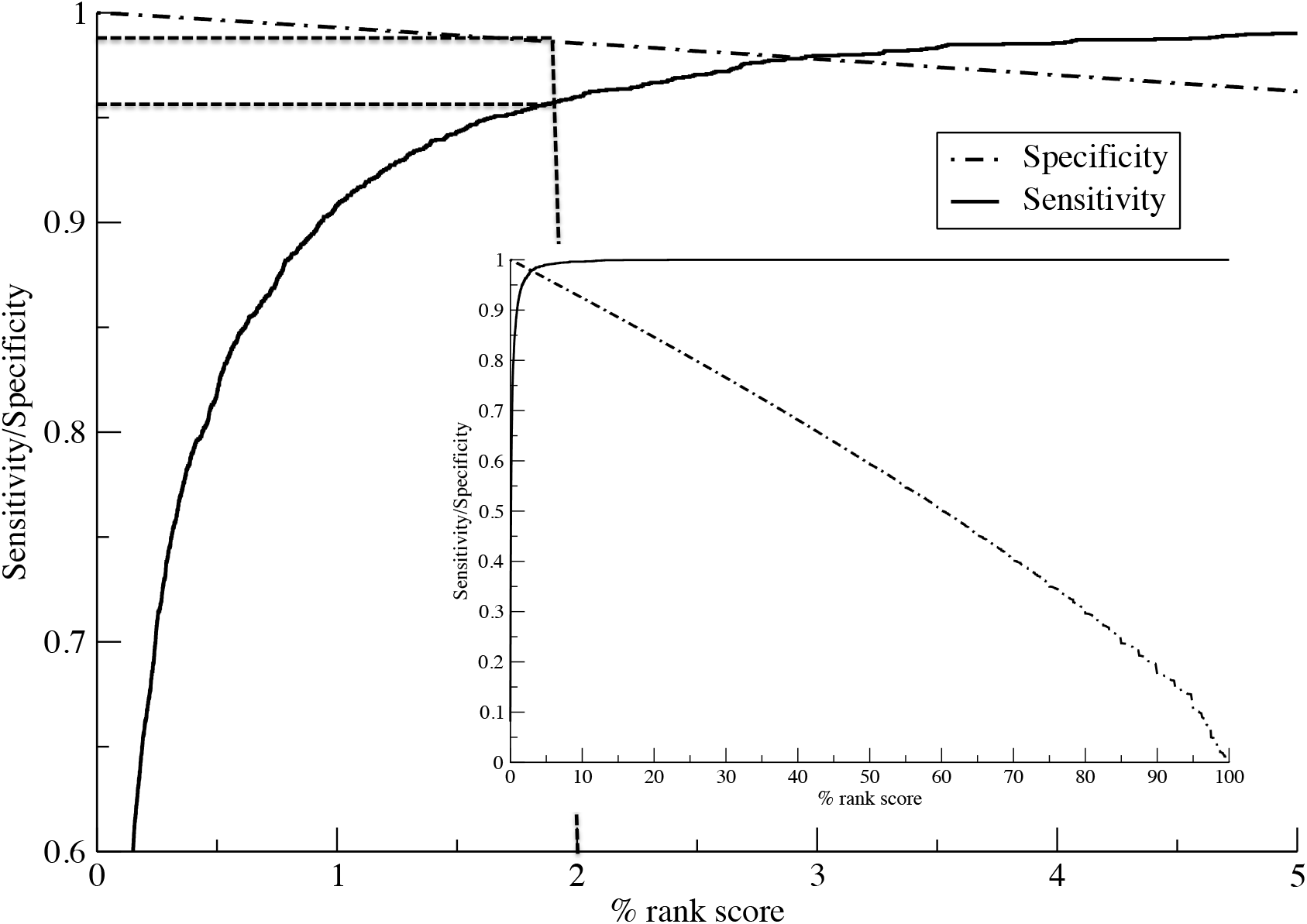
Sensitivity and specificity performance curves for the NetMHCpan-4.0 eluted ligand likelihood predictions. Curves are estimated from a balanced set of eluted ligands from the (17) data set. The insert shows the complete sensitivity and specificity curves as a function of the percentile rank score. The main plot shows the curves in the high-scoring range for 0-5 percentile scores. Dotted vertical and horizontal lines are guides to the eye indicating sensitivity and specificity and the 2% rank score threshold.

### Evaluation on unbiased data sets

Most eluted ligand data potentially suffer from biases towards current prediction methods. This is because many eluted ligand studies, including the Pearson data used here, assign MHC restriction based on predicted binding. To address the impact of this bias, we here benchmark our method against sets of unfiltered eluted ligand data. These data sets were obtained from Bassani-Sternberg et al. (22), and cover eluted ligands obtained from 6 cell lines each with typed HLA expression. From these data, we constructed 6 benchmark data sets by enriching each positive eluted ligand data set with a set of random natural negative peptides (for details see materials and methods). After filtering out data included in the training data of NetMHCpan-4.0, we next benchmarked the predictive power of the different prediction methods. The result of the benchmark is shown in figure 9.

**Figure 9:**
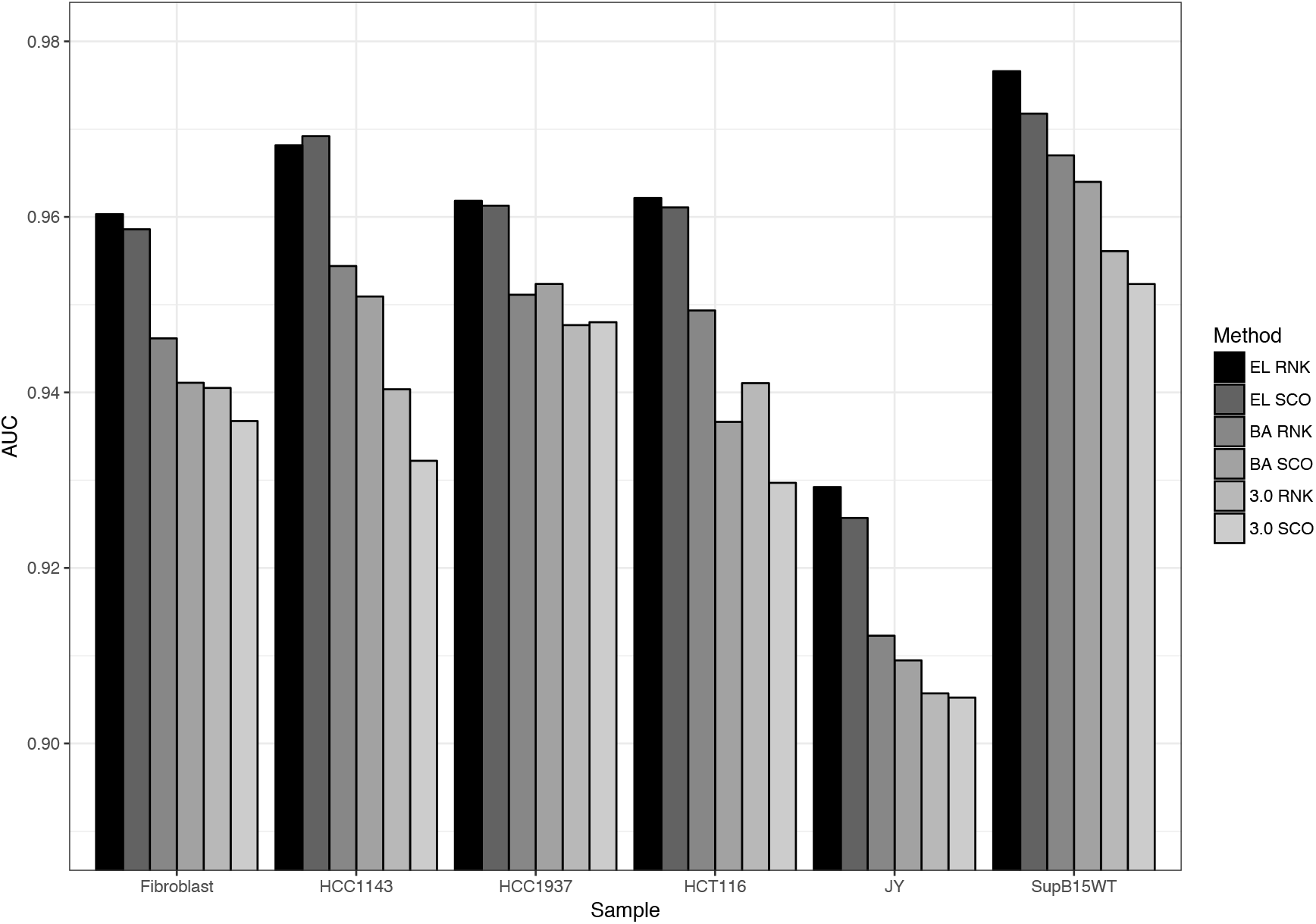
Predictive performance measured in terms of AUC on the Bassani-Sternberg unfiltered eluted ligand data sets. Prediction values are assigned to each peptide in a given data set as the lowest percentile rank score / highest prediction score to each of the HLA molecule expressed by the given cell line. The six methods included are: EL RNK (NetMHCpan-4.0 eluted ligand percentile rank), EL SCO (NetMHCpan-4.0 eluted ligand likelihood score), BA RNK (NetMHCpan-4.0 binding affinity percentile rank), BA SCO (NetMHCpan-4.0 binding affinity score), 3.0 RNK (NetMHCpan-3.0 percentile rank, and 3.0 SCO (NetMHCpan-3.0 binding affinity score).

These results clearly confirm the improved performance of the proposed NetMHCpan-4.0 eluted ligand likelihood predictions over both the NetMHCpan-4.0 and NetMHCpan-3.0 binding affinity predictions. Also, the results show that in the majority of cases the percentile rank predictions achieve improved predictive performance compared to the raw prediction scores.

### Identification of cancer neoantigens

A research field where prediction of naturally processed and presented eluted ligand has attracted large recent attention is rational identification of cancer neoantigens. In contrast to tumor-associated self-antigens, cancer neoantigen are naturally presented ligands containing tumour-specific mutations. Such neoantigens are attracting large attention since these peptides are new to the immune system and not found in normal tissues, and hence are ideal potential cancer vaccine candidates or targets for adoptive T cell therapy. Depending on the mutational load, the number of potential tumour-specific neopeptides (peptides containing one or more missense mutations) can be in the order of many thousands (25). This large number of potential peptide candidates clearly underlines the need for tools to rationally downsize the peptide space in the search for cancer neoepitopes. A recent study by Bassani-Sternberg et al. (14) demonstrated how this downsizing could be effectively achieved by a prediction method trained on a large set of MS eluted ligands. Here, we repeated this benchmark analysis using NetMHCpan-4.0. The results are shown in figure 10 and confirm the finding by Bassani-Sternberg et al. (14), that predictors trained on MS eluted ligand data information in most cases show very high predictive power for the identification of cancer neoantigens. Both the NetMHCpan-4.0 and MixMHCpred method proposed by Bassani-Sternberg et al. (14) identify the known neoantigens within the top 25 peptides in 6 out out 10 cases. NetMHCpan-3.0 only achieves this in 2 out of 10 cases. The results also confirm the earlier findings presented here, that NetMHCpan-4.0 achieves improved performance compared to that of version 3.0, and that the ligands in all cases are predicted with very strong eluted ligand likelihood values (all percentile rank values are less than 1, and the majority are less than or equal to 0.02).

**Figure 10:**
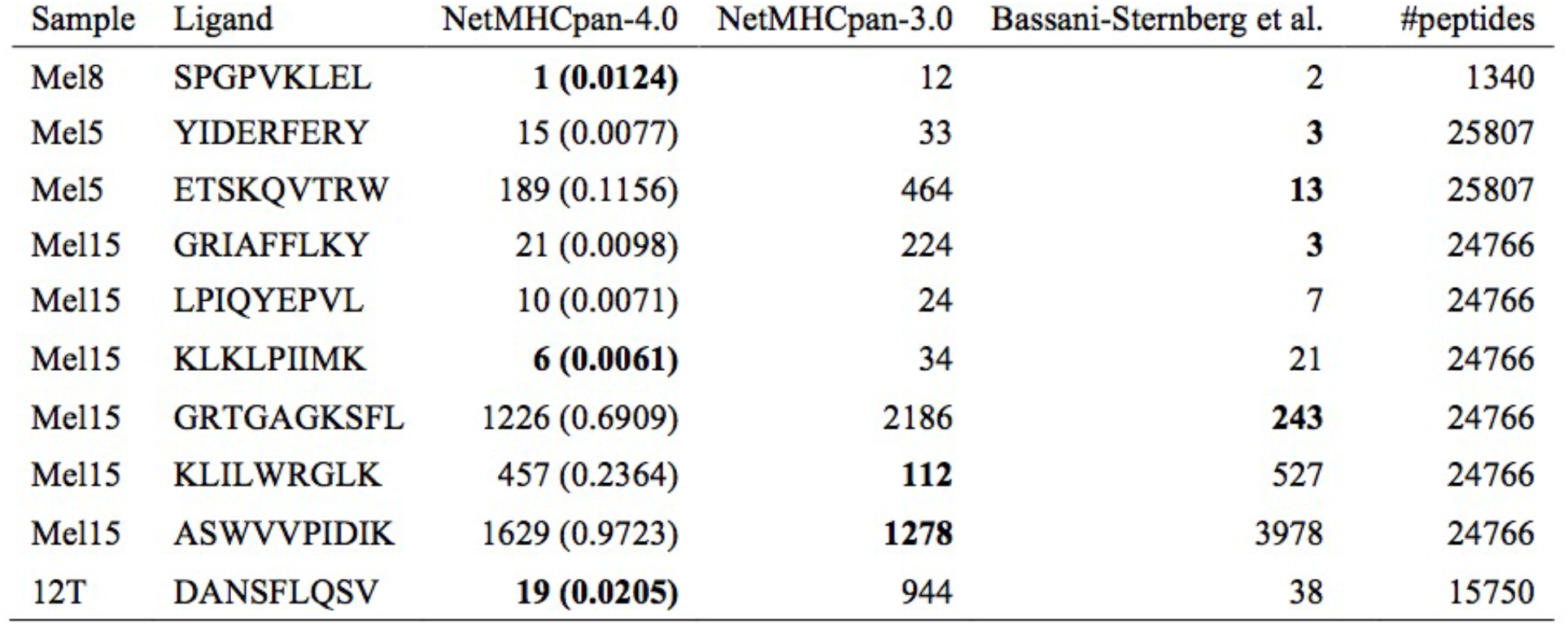
Predictive performance evaluated in terms of rank of neo-antigens identified in four melanoma samples. A rank value of 1 corresponds to the ligand obtaining the highest score (lowest percentile rank) of all peptides from the given sample. Data and performance values for MixMHCFpred are from (31). NetMHCpan-4.0 and NetMHCpan-3.0 are performance values obtained by assigning to each peptide in the given data set the lowest percentile rank score to each of the HLA-A and B molecules expressed by the given cell line. The values in parentheses for NetMHCpan-4.0 are the predicted percentile rank values. Lowest rank value for each ligand is highlighted in bold.

## Discussion

In this work, we have demonstrated how a relatively simple pan-specific machine learning method based on the NNAlign framework can be constructed integrating information from binding affinity data with MS peptidome data. Benefitting from the larger set of peptide binding affinity data with very broad MHC coverage (more than 150 molecules), and the additional information contained within MS peptideome data (information about both antigen processing and presentation, and allele specific peptide length profile), we could demonstrate that the proposed method, NetMHCpan-4.0, achieved improved predictive performance not only when it comes to characterizing the binding specificity of a given MHC molecule, but also when it comes to predicting the peptide length profile. Due to the pan-specific potential of the method, the improved performance was extended beyond the relatively few MHC molecules characterized by MS binding data included in the training of the method. Given this, we thus conclude that the proposed framework is able to benefit from the best of the two data sets; MHC coverage from the binding affinity data, and antigen processing and presentation, and allele specific peptide length profile from the MS data.

Our benchmarks confirmed earlier findings that prediction values for known ligands vary substantially between MHC molecules (26), and that only by the use of percentile rank scores can predictions between different MHC molecules be readily compared.

The improved peptide-MHC tool is made publicly available at www.cbs.dtu.dk/services/NetMHCpan-4.0. The tool was benchmarked on two large independent data sets; one consisting of ~16,000 MS identified MHC restricted ligands (17) and one consisting of more than 1,250 validated T cell epitopes described in the IEDB. For both data sets, the updated version 4.0 of NetMHCpan significantly outperformed the earlier NetMHCpan 3.0 method. In particular, the benchmark on T cell epitope data - to the best of our knowledge - demonstrated for the first time how integration of MS peptidome data into a prediction method of MHC peptide presentation, can lead to improved predictive performance for T cell epitope discovery. The improved performance on this data set was only observed for the method trained on the combined data, and was not observed for the method trained on MS peptidome data alone. This observation underlines the large benefit of merging the two data types.

Investigating potential causes for the observed improved performance of the proposed tool for identification of eluted ligands confirmed earlier findings that eluted ligands share a reduced amino acid diversity at the MHC anchor positions (13). This observation is consistent with the notion that ligands are more stably bound to MHC-I molecules compared with average affinity-defined bound peptides. We postulate that this difference in binding preferences between eluted ligand and affinity-defined peptide binders, combined with the improved prediction accuracy of ligand length preference of the EL methods are the main factors driving the improved predictive performance.

When benchmarking the predictive performance for identification of T cell epitopes, we observed that only the NetMHCpan-4.0 EL model trained on the combined eluted ligand and binding affinity data set demonstrated an improved predictive performance compared to NetMHCpan-3.0. This observation was surprising at first, as we would expect an improved performance also by the method trained on the eluted ligand only due to the reasons outlined above. One likely explanation for this result is the bias in the T cell epitope data towards predicted binding affinity motifs. Most T cell epitopes have been identified using some kind of HLA binding predictions as a filter prior to experimental validation hence giving a bias towards prediction methods trained based on binding affinity data. Given this, the source of the improved performance of the NetMHCpan-4.0 EL method compared to NetMHCpan-4.0 BA on the T cell epitope benchmark data set is thus primarily driven by its improved prediction of the ligand length preference.

It is clear that even with the improved predictive performance of the NetMHCpan-4.0 tool reported here, not all MHC ligands and T cell epitopes will be captured by a prediction workflow. Likewise, it is clear that very few if any experimental workflows enable the exhaustive identification of the ligandome or epitope set contained within a given sample. Given the two workflows to work in concert and use *in-silico* screens as a guide to the experimental setup to effectively boost the sensitivity of the combined workflow. Such an approach where *in-silico* predictions were used to reduce the search space has with success been used to improve the sensitivity of MHC class I ligand discovery (27) and we expect other similar applications to appear in the future.

The machine-learning framework proposed here is not limited to the integration of MHC class I peptide binding affinity and MS peptidome data. The approach can readily be extended to integrate other types of relevant data including MHC binding stability (28), and epitope data. Also, the approach can in its current form be directly applied to the MHC class II system. The only critical limitation for such data integrations is the criteria that each data point must be associated with a specific MHC element. This information is not always readily available, but can in most cases be inferred by unsupervised clustering of the available data (using GibbsCluster (29), position weight matrix mixture models (16), or similar approaches), and association of each cluster to an MHC molecule of the given host.

In conclusion, we have here described a new framework for training of prediction methods for MHC peptide presentation prediction integrating information from two data sources (MS eluted ligand and peptide binding affinity). The framework was used to develop an updated version of NetMHCpan (version 4.0, available at www.cbs.dtu.dk/services/NetMHCpan-4.0) with improved predictive performance for identification of validated eluted ligands, cancer neoantigens and T cell epitopes.

## References

1. Nielsen, M., and M. Andreatta. 2016. NetMHCpan-3.0; improved prediction of binding to MHC class I molecules integrating information from multiple receptor and peptide length datasets. Genome Med. 8: 1–9.

2. Vita, R., J. A. Overton, J. A. Greenbaum, J. Ponomarenko, J. D. Clark, J. R. Cantrell, D. K. Wheeler, J. L. Gabbard, D. Hix, A. Sette, and B. Peters. 2015. The immune epitope database (IEDB) 3.0. Nucleic Acids Res 43: D405–12.

3. Nielsen, M., and M. Andreatta. 2017. NNAlign: a platform to construct and evaluate artificial neural network models of receptor-ligand interactions. Nucleic Acids Res..

4. Andreatta, M., and M. Nielsen. 2015. Gapped sequence alignment using artificial neural networks: application to the MHC class I system. Bioinformatics.

5. Deres, K., T. N. Schumacher, K. H. Wiesmuller, S. Stevanovic, G. Greiner, G. Jung, and H. L. Ploegh. 1992. Preferred size of peptides that bind to H-2 Kb is sequence dependent. Eur J Immunol 22: 1603–8.

6. Trolle, T., C. P. McMurtrey, J. Sidney, W. Bardet, S. C. Osborn, T. Kaever, A. Sette, W. H. Hildebrand, M. Nielsen, and B. Peters. 2016. The Length Distribution of Class I-Restricted T Cell Epitopes Is Determined by Both Peptide Supply and MHC Allele-Specific Binding Preference. J Immunol 196: 1480–7.

7. Lundegaard, C., K. Lamberth, M. Harndahl, S. Buus, O. Lund, and M. Nielsen. 2008. NetMHC-3.0: accurate web accessible predictions of human, mouse and monkey MHC class I affinities for peptides of length 8-11. Nucleic Acids Res 36.

8. Nielsen, M., C. Lundegaard, T. Blicher, K. Lamberth, M. Harndahl, S. Justesen, G. Roder, B. Peters, A. Sette, O. Lund, and S. Buus. 2007. NetMHCpan, a method for quantitative predictions of peptide binding to any HLA-A and-B locus protein of known sequence. PLoS ONE 2: e796.

9. Kreiter, S., M. Vormehr, N. van de Roemer, M. Diken, M. Löwer, J. Diekmann, S. Boegel, B. Schrörs, F. Vascotto, J. C. Castle, A. D. Tadmor, S. P. Schoenberger, C. Huber, Ö. Türeci, and U. Sahin. 2015. Mutant MHC class II epitopes drive therapeutic immune responses to cancer. Nature 520: 692–696.

10. Gubin, M. M., M. N. Artyomov, E. R. Mardis, and R. D. Schreiber. 2015. Tumor neoantigens: building a framework for personalized cancer immunotherapy. J. Clin. Invest. 125: 3413–3421.

11. 2017. The problem with neoantigen prediction. Nat. Biotechnol. 35: 97.

12. Tenzer, S., B. Peters, S. Bulik, O. Schoor, C. Lemmel, M. M. Schatz, P. M. Kloetzel, H. G. Rammensee, H. Schild, and H. G. Holzhutter. 2005. Modeling the MHC class I pathway by combining predictions of proteasomal cleavage, TAP transport and MHC class I binding. Cell Mol Life Sci 62: 1025–37.

13. Harndahl, M., M. Rasmussen, G. Roder, I. Dalgaard Pedersen, M. Sorensen, M. Nielsen, and S. Buus. 2012. Peptide-MHC class I stability is a better predictor than peptide affinity of CTL immunogenicity. Eur J Immunol 42: 1405–16.

14. Bassani-Sternberg, M., C. Chong, P. Guillaume, M. Solleder, H. Pak, P. O. Gannon, L. E. Kandalaft, G. Coukos, and D. Gfeller. 2017. Deciphering HLA-I motifs across HLA peptidomes improves neoantigen predictions and identifies allostery regulating HLA specificity. PLOS Comput. Biol. 13: e1005725.

15. Abelin, J. G., D. B. Keskin, S. Sarkizova, C. R. Hartigan, W. Zhang, J. Sidney, J. Stevens, W. Lane, G. L. Zhang, T. M. Eisenhaure, K. R. Clauser, N. Hacohen, M. S. Rooney, S. A. Carr, and C. J. Wu. 2017. Mass Spectrometry Profiling of HLA-Associated Peptidomes in Mono-allelic Cells Enables More Accurate Epitope Prediction. Immunity 46: 315–326.

16. Bassani-Sternberg, M., and D. Gfeller. 2016. Unsupervised HLA Peptidome Deconvolution Improves Ligand Prediction Accuracy and Predicts Cooperative Effects in Peptide-HLA Interactions. J. Immunol. Baltim. Md 1950 197: 2492–2499.

17. Pearson, H., T. Daouda, D. P. Granados, C. Durette, E. Bonneil, M. Courcelles, A. Rodenbrock, J.-P. Laverdure, C. Côté, S. Mader, S. Lemieux, P. Thibault, and C. Perreault. 2016. MHC class I-associated peptides derive from selective regions of the human genome. J. Clin. Invest. 126: 4690–4701.

18. Bassani-Sternberg, M., E. Bräunlein, R. Klar, T. Engleitner, P. Sinitcyn, S. Audehm, M. Straub, J. Weber, J. Slotta-Huspenina, K. Specht, M. E. Martignoni, A. Werner, R. Hein, D. H Busch, C. Peschel, R. Rad, J. Cox, M. Mann, and A. M. Krackhardt. 2016. Direct identification of clinically relevant neoepitopes presented on native human melanoma tissue by mass spectrometry. Nat. Commun. 7: 13404.

19. Sidney, J., S. Southwood, C. Oseroff, M. F. del Guercio, A. Sette, and H. M. Grey. 2001. Measurement of MHC/peptide interactions by gel filtration. Curr. Protoc. Immunol. Chapter 18: Unit 18.3.

20. Harndahl, M., S. Justesen, K. Lamberth, G. Roder, M. Nielsen, and S. Buus. 2009. Peptide binding to HLA class I molecules: homogenous, high-throughput screening, and affinity assays. J. Biomol. Screen. 14: 173–80.

21. Nielsen, M., C. Lundegaard, P. Worning, S. L. Lauemoller, K. Lamberth, S. Buus, S. Brunak, and O. Lund. 2003. Reliable prediction of T-cell epitopes using neural networks with novel sequence representations. Protein Sci 12: 1007–17.

22. Bassani-Sternberg, M., S. Pletscher-Frankild, L. J. Jensen, and M. Mann. 2015. Mass spectrometry of human leukocyte antigen class I peptidomes reveals strong effects of protein abundance and turnover on antigen presentation. Mol Cell Proteomics 14: 658–73.

23. Jorgensen, K. W., M. Rasmussen, S. Buus, and M. Nielsen. 2013. NetMHCstab - predicting stability of peptide:MHC-I complexes; impacts for CTL epitope discovery. Immunology.

24. Paul, S., D. Weiskopf, M. A. Angelo, J. Sidney, B. Peters, and A. Sette. 2013. HLA class I alleles are associated with peptide-binding repertoires of different size, affinity, and immunogenicity. J Immunol 191: 5831–9.

25. Bjerregaard, A.-M., M. Nielsen, S. R. Hadrup, Z. Szallasi, and A. C. Eklund. 2017. MuPeXI: prediction of neo-epitopes from tumor sequencing data. Cancer Immunol. Immunother. CII.

26. Karosiene, E., M. Rasmussen, T. Blicher, O. Lund, S. Buus, and M. Nielsen. 2013. NetMHCIIpan-3.0, a common pan-specific MHC class II prediction method including all three human MHC class II isotypes, HLA-DR, HLA-DP and HLA-DQ. Immunogenetics 65: 711–24.

27. Murphy, J. P., P. Konda, D. J. Kowalewski, H. Schuster, D. Clements, Y. Kim, A. M. Cohen, T. Sharif, M. Nielsen, S. Stevanovic, P. W. Lee, and S. Gujar. 2017. MHC-I Ligand Discovery Using Targeted Database Searches of Mass Spectrometry Data: Implications for T-Cell Immunotherapies. J. Proteome Res. 16: 1806–1816.

28. Rasmussen, M., E. Fenoy, M. Harndahl, A. B. Kristensen, I. K. Nielsen, M. Nielsen, and S. Buus. 2016. Pan-Specific Prediction of Peptide-MHC Class I Complex Stability, a Correlate of T Cell Immunogenicity. J Immunol 197: 1517–24.

29. Andreatta, M., B. Alvarez, and M. Nielsen. 2017. GibbsCluster: unsupervised clustering and alignment of peptide sequences. Nucleic Acids Res..

30. Thomsen, M. C., and M. Nielsen. 2012. Seq2Logo: a method for construction and visualization of amino acid binding motifs and sequence profiles including sequence weighting, pseudo counts and twosided representation of amino acid enrichment and depletion. Nucleic Acids Res 40: W281–7.

31. Bassani-Sternberg, M., C. Chong, P. Guillaume, M. Solleder, H. Pak, P. O. Gannon, L. E. Kandalaft, G. Coukos, and D. Gfeller. 2017. Deciphering HLA-I motifs across HLA peptidomes improves neoantigen predictions and identifies allostery regulating HLA specificity. bioRxiv 98780.

